# Gaining insight into the non-focality of beta oscillation suppression along the sensorimotor cortex using corticomuscular coherence

**DOI:** 10.64898/2026.01.15.699672

**Authors:** Christian Georgiev, Scott J. Mongold, Gilles Naeije, Mathieu Bourguignon

## Abstract

An Event-Related Desynchronization (ERD) of the 13–30 Hz sensorimotor (SM1) beta oscillations is commonly observed during movement preparation and execution. Human electrophysiological measurements suggest that such a beta ERD has a wide topographical distribution along the SM1, however, no accessible means of quantifying the degree of its focality exist. Here, we tested the suitability of a method to investigate how the movement-induced beta ERD in one somatotopic SM1 area affects beta oscillations in a neighbouring SM1 area. Thirty-six participants performed right brachium movements while holding a submaximal isometric contraction with their right first dorsal interosseous (FDI) muscle. Beta ERD in the left SM1 brachium area was assessed with electroencephalography (EEG). The effect of that ERD on beta activity in the neighbouring SM1 FDI area was assessed through the corticomuscular coherence (CMC) between SM1 EEG signals and the electromyography and force signals recorded from the stationary isometrically contracted right FDI muscle. Our results showed a strong movement-induced beta ERD in the SM1 brachium area that co-occurred with an attenuation of CMC with both FDI signals. These findings imply that beta ERD may not be a strictly focal phenomenon as it could spread to a neighbouring SM1 area. Importantly, we introduced a novel approach that combines a dual motor task paradigm with CMC to assess beta propagatory effects. This approach could allow for investigating the topographical properties of beta oscillations and their role for motor control in healthy and clinical populations.

## 1. Introduction

Synchronous activity of primary sensorimotor cortical (SM1) pyramidal neurons gives rise to a bursty oscillation at ∼20 Hz, commonly referred to as the beta rhythm and typically captured with electroencephalography (EEG) and magnetoencephalography (MEG; Démas et al., 2020; Engel & Fries, 2010; Kilavik et al., 2013). The ∼100 ms bursts which comprise the beta oscillations have a functional significance for motor behavior, with a suppression of their amplitude and rate occurring shortly before and during bodily movement and an enhancement of their amplitude and rate occurring after movement (Georgiev et al., 2025; Heinrichs-Graham et al., 2017; Khanna & Carmena, 2017; Shin et al., 2017; Szul et al., 2023; Wessel, 2020). These patterns of modulation can be seen in a trial-averaged time-frequency representation (TFR), where global beta amplitude suppression and enhancement are classically termed Event-Related Desynchronization (ERD) and Event-Related Synchronization (ERS), respectively (Nakayashiki et al., 2014; Pfurtscheller, 1981; Salenius & Hari, 2003; Zaepffel et al., 2013).

Investigations of the beta oscillations during movement execution have consistently identified the neural sources of beta ERD to be located predominantly in the SM1 contralateral to the moving limb (Bonaiuto et al., 2021; Murthy & Fetz, 1996; Tzagarakis et al., 2010; Zich et al., 2023). A degree of focality of beta oscillations within the SM1 homunculus has been shown, with maximal beta ERD observed above the somatotopic area governing each given contracting contralateral muscle (Bonaiuto et al., 2021; Shin et al., 2017; Stancák et al., 2000). However, how such focal beta suppression in one somatotopic area affects the beta oscillatory activity of neighbouring somatotopic areas remains unclear. Typically, EEG and MEG scalp topography and source reconstruction of beta ERD appear to be distributed along the SM1, spanning several somatotopic areas (Bourguignon et al., 2017; Hari et al., 2014). Furthermore, evidence from local field potential recordings in primates (Balasubramanian et al., 2020; Best et al., 2017) as well as MEG recordings in humans (Zich et al., 2023) has identified a spatio-temporal propagation of beta burst suppression along the SM1 (Balasubramanian et al., 2020; Best et al., 2017). This wider beta suppression could play a role in priming potential neural activation of different SM1 areas and hence preparing for the execution of more complex bodily movements and their combination into purposive actions. Characterizing the focality of beta ERD empirically, however, is not straightforward due to certain shortcomings of non-invasive human electrophysiology. Due to field spread, EEG and MEG sensors detect activity from multiple somatotopic areas (Hämäläinen et al., 1993). Moreover, due to the ill-posed nature of the inverse problem (Grech et al., 2008), EEG and MEG source reconstruction techniques do not allow for precise distinction between anatomically adjacent sources.

The aim of the present work was to characterize the focality of the beta oscillations along the SM1 by investigating how the movement-induced beta ERD and subsequent ERS in one somatotopic area affects beta oscillatory activity in neighbouring areas. To circumvent the limitations outlined above, we introduced a novel approach for investigating the impact of beta amplitude changes in one SM1 area on the corticomuscular coherence (CMC) between a neighbouring SM1 area and its target muscle. CMC is a technique that quantifies the coupling between SM1 signals and the electromyography (EMG) of a contralateral isometrically contracted muscle (Bourguignon et al., 2019). CMC typically peaks at ∼20 Hz beta frequency (Bourguignon et al., 2017; Conway et al., 1995; Kilner et al., 2000; Mendez-Balbuena et al., 2012; Piitulainen et al., 2015; Steeg et al., 2014) and originates from the SM1 somatotopic representation of the contracted muscle (Bourguignon et al., 2017; Brown et al., 1998; Maezawa et al., 2014). Furthermore, CMC is contingent on the presence of high-amplitude ∼20 Hz beta bursts in that somatotopic SM1 area that are forwarded to the contracted muscle (Bourguignon et al., 2017; Echeverria-Altuna et al., 2022; Mongold et al., 2022). Therefore, the CMC between a brain area and a target muscle could be used as a proxy for the amplitude dynamics of the beta oscillations within that given brain area.

Following this rationale, we conducted an experiment with a dual movement execution task. During the task, participants were prompted to perform small transient internal/external rotations of the right brachium. Concurrently, the CMC between the left SM1 and the stationary isometrically contracted right first dorsal interosseous (FDI) muscle was recorded. This setup capitalizes on the distinct, yet closely neighbouring, SM1 somatotopic representations of the brachium muscles (subscapularis, teres minor, and infraspinatus) and the FDI. According to classical views on somatotopic organization (Penfield & Boldrey, 1937), the brachium and FDI SM1 areas should be distinct from one another and should have distinct intrinsic beta oscillatory patterns (Neuper & Pfurtscheller, 2001). In an EEG experiment where each sensor detects a mixture of activity from multiple underlying SM1 areas, if brachium movement-induced beta ERD is focal, the beta ERD in the SM1 brachium area should allow for enhanced detection of the beta oscillations from the SM1 FDI area. This is expected because beta oscillations from the brachium area and those from the FDI area both contribute to the overall signal detected at the given EEG electrode (alongside other contributions unrelated to the task). Hence, the transient suppression of the beta oscillations from the SM1 brachium area would leave the unsuppressed beta oscillations from the FDI area more salient in the EEG signal and thus result in enhanced CMC between that area and the FDI muscle. Alternatively, if movement-induced beta ERD is non-focal, executing the rotational brachium movement should result in a beta ERD in the SM1 brachium area which spreads to the SM1 FDI area. Hence, transient suppression of the beta oscillations from both SM1 areas would globally suppress all beta activity in the EEG signal and thus attenuate the CMC between that FDI area and the FDI muscle. The application of such a CMC-based characterization of non-focal beta oscillatory activity would allow for the development of approaches for non-invasively quantifying the spread of beta oscillations along the SM1 homunculus in healthy and clinical populations.

## 2. Methods

### 2.1 Participants

Thirty-six adults (15 female; Mean ± SD age = 25 ± 4 years) participated in the study. All participants were right-handed (Mean ± SD Edinburgh Handedness Inventory = 88 ± 14), did not suffer from neurological or psychiatric disorders, and had normal or corrected-to-normal vision and hearing. The study protocol was approved by the local ethics committee (Comité d’Ethique Hospitalo-Facultaire Erasme-ULB, 021/406, Brussels, Belgium) and all participants gave written informed consent prior to participating.

### 2.2 Experimental Protocol

Participants’ FDI maximum voluntary contraction (MVC) force was measured as the mean force applied during a 5 s maximal pinch grip contraction with the right thumb and index finger against a rigid force transducer (CS 50-3Q1, SAUTER GmbH, Wutöschingen, Germany). The participants were then seated on a chair with their arms placed on the armrests. Their task was to maintain a submaximal isometric pinch grip contraction with their right thumb and index finger against a custom-build force transducer with a 6 N load cell (Vishay Precision Group, Malvern, PA, USA) at 10 ± 3 % MVC (Figure 1A). While maintaining the contraction, an auditory tone (presented once every 4 s ± 0.5 s random jitter) prompted the participants to perform a 10° internal/external rotation of the right brachium, which resulted in a small lateral movement of the contracted lower arm in the horizontal plane. To facilitate task performance, participants were presented with real-time visual feedback which indicated when the applied force was outside of the 10 ± 3 % MVC range (Figure 1B). This dual movement execution task was performed for two blocks of 5 minutes, separated by a short break.

**Figure 1.**
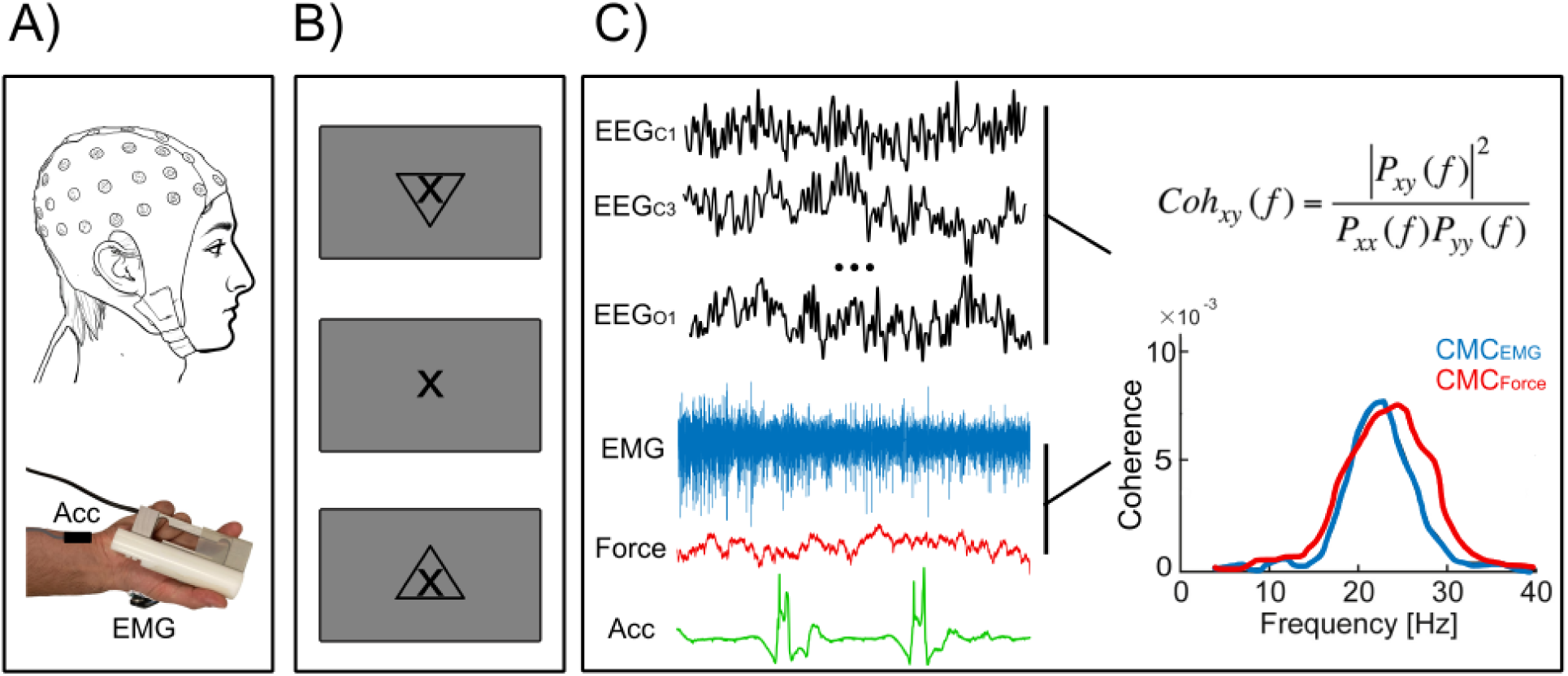
Experimental setup. A — Illustration of the experimental protocol, during which EEG, EMG, force, and accelerometry (Acc) data was acquired while participants performed a dual movement execution and FDI contraction task. B — Visual feedback of the applied force presented to the participants during the dual task. The presentation of an “X” indicated that the force was within the desired range; the presentation of an arrow pointing up indicated that the applied force should increase; and the presentation of an arrow pointing down indicated that the applied force should decrease. C — An illustration of the recorded signals and the procedure for computing CMC.

During the task, 64-channel EEG (EEGO Mylab, ANT Neuro, Hengelo, the Netherlands) was recorded. Prior to recording, electrode impedance was brought below 20 kΩ via the application of electroconductive gel. The EEG recording was online referenced to electrode CPz and sampled at 1000 Hz. Also recorded were: 1) the activity of the contracted right FDI muscle via surface EMG (COMETA Pico, COMETA, Milan, Italy; with an active electrode on the FDI muscle belly and a reference electrode on the first metacarpal bone), 2) the contraction force via the 6 N custom-build force transducer (Vishay Precision Group, Malvern, PA, USA), and 3) the signals of a 3-axis accelerometer (ADXL335, AnalogDevices, Wilmington, MA, USA) attached to the wrist of the moving right hand. All peripheral signals were sampled at 1000 Hz and synchronized with the EEG recordings via the delivery of digital triggers.

### 2.3 Data preprocessing

2.3.1 ***EEG preprocessing***

EEG signals were preprocessed with custom-made Matlab (Mathworks, Natick, MA) scripts along with functions from FieldTrip (Oostenveld et al., 2011). EEG signals at electrodes affected by excessive noise levels were reconstructed by topographic interpolation. Electrodes were considered excessively noisy when they had: 1) excessive wide-band amplitude, 2) excessive ratio between high- and low-frequency amplitudes, and 3) a low correlation with neighbouring channels. Further details about this approach can be found in Bigdely-Shamlo et al. (2015). Subsequently, EEG signals were re-referenced to the common average and filtered through 0.3–45 Hz. Independent Components Analysis (Vigário et al., 2000) was used for further artifact suppression. Twenty independent components were evaluated from the data with a FastICA algorithm with dimension reduction, 25 nonlinearity, tanh (Hyvärinen & Oja, 2000; Vigário et al., 2000) and independent components corresponding to eye-blink, eye movement, and heartbeat artifacts were removed. On average, 4 (SD = 1) independent components per participant were removed. Finally, time points where full-band EEG amplitude in at least one electrode was greater than 5 SDs above the mean were set to bad.

#### 2.3.2 EMG and force preprocessing

The FDI EMG was not filtered and not rectified, in accordance with the methodology of previous successful experiments from our lab (Bourguignon et al., 2017; Hari et al., 2014; Mongold et al., 2022; Piitulainen et al., 2015). Time points where the applied force was outside of the 10 ± 3 % MVC were marked as bad and removed from all data.

### 2.4 Data Analyses

#### 2.4.1 Trial extraction

Continuous EEG data were filtered through 5-Hz-wide frequency bands centered on 5–40 Hz by 1 Hz steps. The applied bandpass filters were designed in the frequency domain with zero-phase and 1-Hz-wide squared-sine transitions from 0 to 1 and 1 to 0 (e.g., the filter centered on 20 Hz rose from 0 at 17 Hz to 1 at 18 Hz and ebbed from 1 at 22 Hz to 0 at 23 Hz). EEG signals were smoothly set to 0 (squared-cosine transition of 1 s) at timings 1 s around bad time points. Subband analytical signals were then obtained with Hilbert transformation. They were then divided by the low-frequency content of their envelope fluctuations, obtained as their envelope low-pass filtered at 0.1 Hz, ensuring signal envelopes fluctuate about 1. The data was subsequently segmented into 3-s epochs, from -1 s to 2 s relative to brachium movement onsets. Movement onsets were identified on a trial-by-trial basis based on the 3-axis accelerometer signals recorded from the moving hand. Therefore, the 3-s epochs were all aligned with respect to movement onset. Epochs which contained time points less than 2 s away from bad time points or in which participants failed to perform a movement were excluded from the analysis (on average, per participant 106 ± 40 artifact-free epochs were retained out of 170 epochs in total).

#### 2.4.2 Beta ERD and ERS analyses

The envelopes of the retained epochs’ analytical signals were subsequently averaged for each participant, giving rise to a time-frequency map (TFR) of amplitude modulation for each EEG electrode. These maps were then converted to relative amplitude change maps by subtracting 1. This subtraction allows for straightforward quantification of the brachium movement-induced changes in amplitude in terms of percentage change relative to baseline (i.e., value 0). All subsequent analyses of the ERD and ERS were performed on EEG electrode C1. This electrode was selected for its location above the right brachium area of the left SM1, where the strongest beta ERD and ERS are expected in response to the transient brachium movements. Moreover, focusing on a single electrode was deemed sufficient for this analysis, since its aim was only to confirm the presence of movement-induced ERD and ERS.

For each participant, we separated the artifact-free epochs into a set with high beta ERD and one with low beta ERD based on a median split. This analysis was based on single epoch ERD values, which were estimated as the mean across 13–30 Hz and - 500 to 800 ms of relative amplitude change maps derived from single epochs.

#### 2.4.3 CMC analyses

Coherence analysis was applied to compute CMC between the signals from all EEG electrodes and the right FDI EMG signal (CMC_EMG_) as well as with the right force signal (CMC_Force_; Figure 1C). Previous MEG studies have demonstrated the equivalence of CMC_EMG_ and CMC_Force_ (Airaksinen et al., 2015; Bourguignon et al., 2017; Piitulainen et al., 2013), however, the force approach in EEG is less common. Therefore, we also set out to compare the fidelity of CMC_Force_ to the more commonly used CMC_EMG_ derived from EEG recordings.

Coherence analyses were performed based on the analytical signals of the EEG, EMG, and force data following the approach described in Bourguignon et al. (2017). CMC_EMG_ and CMC_Force_ were first assessed for each participant globally, by pooling the 3 s of data from all retained epochs. For each participant and type of CMC (CMC_EMG_ and CMC_Force_), the EEG electrode with the greatest global CMC value in the 10–30 Hz band was selected from a subset of electrodes neighbouring to C1 and spanning the SM1 somatotopy, including the area governing the right FDI (FCz, FC1, FC3, FC5, Cz, C3, C5, CP1, CP3, CP5). For further analyses, we retained for each participant the maximum global CMC value at the selected electrode across 13–30 Hz. In addition, we estimated CMC_EMG_ and CMC_Force_ at that electrode, this time separately within three time periods of interest: before the brachium movement (T_before_, from -1 to 0 s), during the brachium movement (T_during_, from 0 to 1 s), and after the brachium movement (T_after_, from 1 to 2 s). These estimates were used to determine whether the movement-induced beta ERD in the right SM1 brachium area affects the CMC with the right FDI. To further clarify this question, the same CMC_EMG_ and CMC_Force_ estimates were also derived from epochs with low and high beta ERD separately. Finally, we estimated the fine-grained time-frequency evolution of CMC_EMG_ and CMC_Force_ at the selected electrode by estimating CMC at every time-frequency bin separately.

#### 2.4.4 FDI EMG and force stability analyses

Differences in CMC_EMG_ and CMC_Force_ across the three time periods of interest were expected to emerge as a result of a non-focal brachium movement-induced beta ERD along the SM1. However, differences in CMC could also emerge as a result of movement-induced, artifact-like changes in FDI EMG and force signals themselves. Hence, to investigate the effect of right brachium movement execution on the stability of the right FDI EMG and force signals, the EMG signal was high-pass filtered at 40 Hz, rectified, and low-pass filtered at 20 Hz and the force signal was band-pass filtered between 2 and 10 Hz. Subsequently, the EMG signal was normalised by its mean and the force signal was normalised by its SD. The two signals were then segmented into 3 s epochs (as done for the EEG data) for each participant with significant CMC (see section **2.5 Statistical Analyses**). Then, for each signal and each segment, the SD in the T_before_, T_during_, and T_after_ time periods was computed as a measure of signal stability.

### 2.5 Statistical Analyses

The statistical significance of relative beta amplitude changes at electrode C1 was assessed with cluster-based non-parametric permutation tests (Maris & Oostenveld, 2007). In brief, the significance of brachium movement-induced modulation in the group-averaged C1 TFR map was investigated by comparing this observed TFR map with 10,000 permutation TFR maps. Each permutation TFR map was obtained by averaging individual TFR maps after having randomly changed the sign of a random selection thereof. Then, for each TFR map, we identified the time-frequency bins within the 13–30 Hz and the -500–1000 ms range where the amplitude was smaller than that in 95% of the other TFR maps. Adjacent bins were then grouped into clusters. The significance of the clusters in the genuine TFR map (i.e., *p*-value) was then obtained as the proportion of clusters in the permutation TFR maps that were larger than them. A similar analysis was conducted for beta ERS within the 13–30 Hz and the 500–2000 ms range. The size of the significant clusters was measured in time-frequency bins (bin size: 50 ms × 1 Hz).

The statistical significance of CMC was evaluated using surrogate data-based statistics as done previously (Mongold et al., 2022). In brief, the distribution of the maximum global CMC — across 13–30 Hz, and across the selection of 10 electrodes — was evaluated between EEG and Fourier-transform surrogate EMG or force signals (1000 repetitions each; Faes et al., 2004). The significance level (i.e., *p*-value) was then obtained as the proportion of values in the surrogate EMG and force distributions that were higher than the genuine ones. Only participants with significant CMC_EMG_ or CMC_Force_ were retained for further analyses.

The impact of the brachium movement on CMC_EMG_ and CMC_Force_ magnitudes, on FDI EMG and force variability (i.e., SD) in the three time periods were assessed with ANOVAs followed by post-hoc *t*-tests. In the case of force variability, the significant ANOVA was also followed by a Spearman correlation analysis between the force SD in the -300 to 500 ms time window (i.e., the period where the movement-induced force variability was greatest) and an index of movement-related CMC_Force_ modulation. Noticing that CMC_Force_ was attenuated during T_before_ and T_during_ compared to T_after_, this index of CMC_Force_ modulation was derived from a contrast between CMC_Force_ averaged across T_before_ and T_during_, and CMC_Force_ at T_after_. More precisely, the index was computed as their difference divided by their sum.

The relationship between the brachium movement-induced beta ERD at electrode C1 and the CMC_EMG_ and CMC_Force_ magnitudes was investigated by comparing the CMC magnitudes between movement execution trials with low beta ERD and high beta ERD with paired-samples *t*-tests. In this comparison, the CMC values being compared were the mean across the T_before_ and T_during_ time windows.

The association and difference between global CMC_Force_ and the more classically reported global CMC_EMG_ were assessed with a Spearman correlation and a paired-samples *t*-test, respectively. We also assessed the differences in the proportion of participants showing a significant CMC_EMG_ and CMC_Force_ with a *χ*^2^ test as well as the consistency in the number of participants showing significant CMC_EMG_ and CMC_Force_ using Cohen’s Kappa test for intercategory association.

## 3. Results

### 3.1 Brachium movement-induced sensorimotor beta ERD

Figure 2 presents the grand-average TFR map at EEG electrode C1. Therein was identified a cluster of significant movement-induced beta ERD (*p* < 0.001) comprising 331 time-frequency bins and spanning the T_before_ and T_during_ time periods. The ERD had an onset ∼500 ms before brachium rotation onset, a maximum suppression at movement onset, and offset at ∼1000 ms after brachium movement onset. Furthermore, the scalp topography of the 13–30 Hz amplitude in the time periods spanning T_before_ and T_during_ revealed a wide pattern of movement-induced beta ERD, compatible with the engagement of the broader arm and hand areas of the left SM1 (Figure 2B). Also noteworthy, is that a cluster of significant alpha (8–12 Hz) ERD comprising 88 time-frequency bins was identified in the T_during_ time period in an a posteriori analysis.

**Figure 2.**
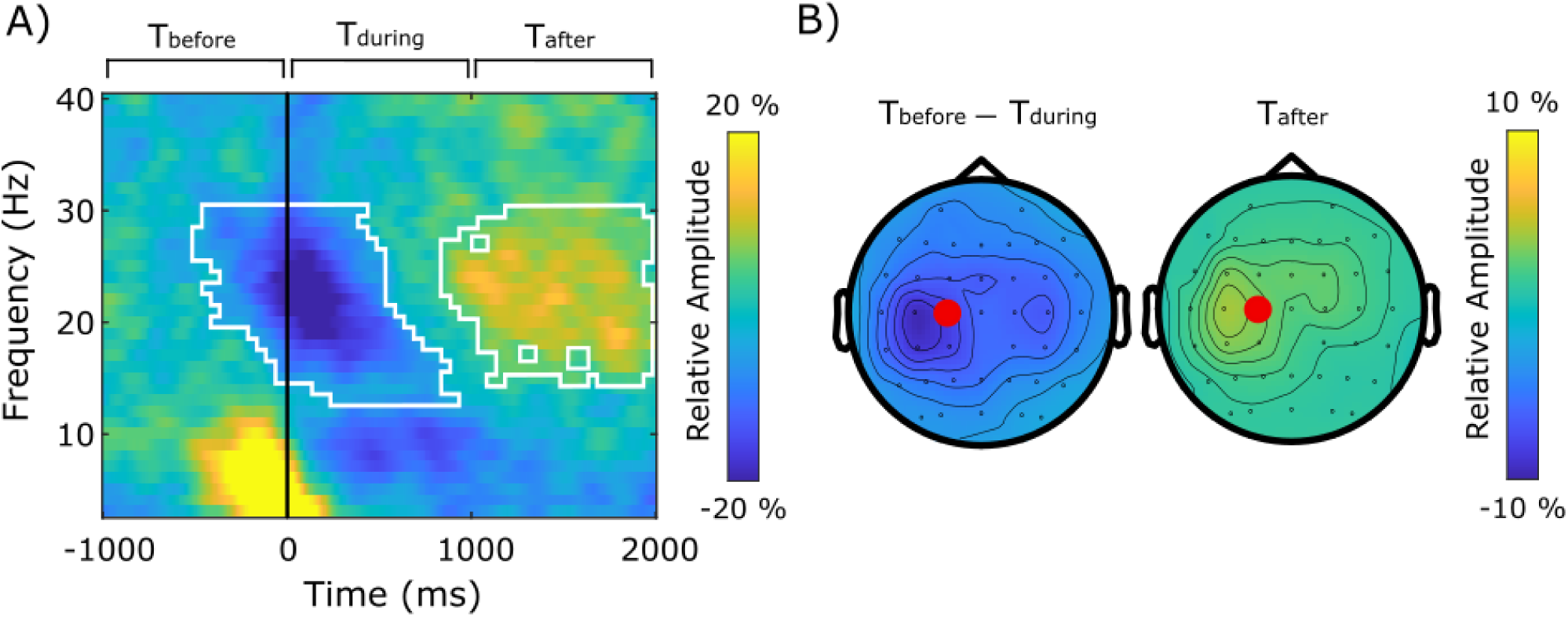
Grand-averaged modulation of EEG relative amplitude in response to the brachium movement execution. A — TFR of EEG at electrode C1. The black vertical line indicates the onset of the brachium rotation as identified in accelerometer recordings. The white lines outline a cluster of statistically significant beta ERD and ERS. B — Scalp distribution of relative EEG amplitude averaged across 13–30 Hz and from -500 to 1000 ms (left) or from 1000 to 2000 ms (right). Red circles denote EEG electrode C1.

In addition to the movement-induced beta ERD, a cluster of significant beta ERS (*p* < 0.001) comprising 307 time-frequency bins was identified in the T_after_ time period. The scalp topography of the 13–30 Hz amplitude in that time period also revealed a wide pattern of beta enhancement compatible with the engagement of the broader arm and hand areas of the left SM1 (Figure 2B).

### 3.2 CMC

Fifteen participants (42 %) displayed significant CMC_EMG_ (0.0001 ≤ *p* ≤ 0.023) and 22 participants (61 %) displayed significant CMC_Force_ (0.0001 ≤ *p* ≤ 0.047). Figure 3 presents the fine-grained time-frequency distribution of CMC_EMG_ and CMC_Force_ across the 3 time periods (Figure 3A), and the averaged values of CMC_EMG_ and CMC_Force_ within the 3 time periods (Figure 3B) as well as their scalp distributions (Figure 3C) for the participants with significant CMC who were retained for further analysis. The scalp distributions highlight a clear CMC peak at SM1 areas neighbouring EEG electrode C1.

**Figure 3.**
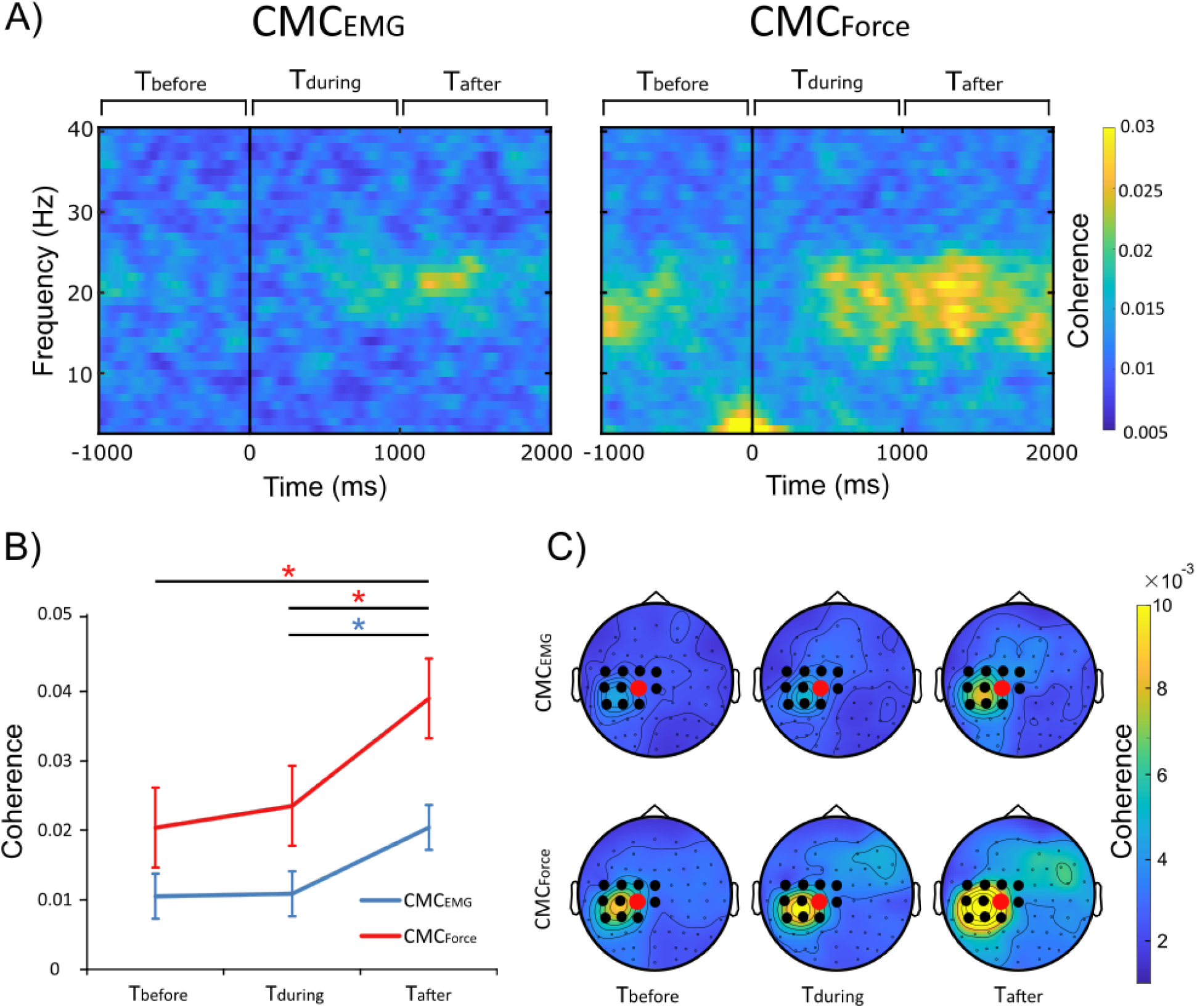
CMC at the three time periods of interest for participants with significant CMC_EMG_ or CMC_Force_. A — Fine-grained time-frequency representation of CMC_EMG_ (left) and CMC_Force_ (right). The black vertical line indicates the onset of the brachium rotation as identified in accelerometer recordings. B — Magnitude of CMC_EMG_ (blue) and CMC_Force_ (red). Error bars depict standard errors of the mean and asterisks indicate significant differences (p < 0.05). C — Scalp distribution of CMC_EMG_ (top) and CMC_Force_ (bottom). Red circles highlight EEG electrode C1, black circles highlight neighbouring electrodes.

Across participants with significant CMC_EMG_, peak CMC_EMG_ frequency was 24.9, 23.4, and 23.3 Hz for the T_before_, T_during_, and T_after_ time periods, respectively, and there was no significant difference in peak frequency between the time periods (*F*_2,42_ = 1.03, *p* = 0.37, *η*^2^ = 0.047). Comparing the magnitude of CMC_EMG_ across T_before_, T_during_, and T_after_ revealed a significant effect of time (*F*_2,28_ = 4.11, *p* = 0.027, *η*^2^ = 0.227) with CMC_EMG_ magnitude being significantly lower at T_during_ compared to T_after_ (*t_14_*= -2.17, *p* = 0.047, *d* = 0.53) and no significant difference in CMC_EMG_ magnitude between T_before_ and T_during_ (*t*_14_ = -0.16, *p* = 0.879, *d* = 0.03), or T_before_ and T_after_ (*t_14_*= -2.12, *p* = 0.052, *d* = 0.53; Figure 3B).

Across participants with significant CMC_Force_, peak CMC_Force_ frequency was 20.5, 20.6, and 20.8 Hz for the T_before_, T_during_, and T_after_ time periods, respectively, and there was no significant difference in peak frequency between the time periods (*F*_2,63_ = 0.03, *p* = 0.97, *η^2^* = 0.001). Comparing the magnitude of CMC_Force_ across T_before_, T_during_, and T_after_ revealed a significant effect of time (*F*_2,42_ = 6.00, *p* = 0.005, *η*^2^ = 0.222) with CMC_Force_ magnitude being significantly lower at T_before_ compared to T_after_ (*t_21_* = -2.69, *p* = 0.014, *d* = 0.38) and at T_during_ compared to T_after_ (*t_21_* = -2.83, *p* = 0.010, *d* = 0.29). There was no significant difference in CMC_Force_ magnitude between T_before_ and T_during_ (*t*_21_ = - 0.68, *p* = 0.50, *d* = 0.08; Figure 3B).

### 3.3 Relationship between brachium movement-induced beta ERD and CMC_EMG_ and CMC_Force_

There was no significant difference in CMC magnitude during T_before_ and T_during_ for brachium movement execution trials with low beta ERD compared to brachium movement execution trials with high beta ERD (CMC_EMG_, *t_14_*= -0.69, *p* = 0.50, *d* = 0.06; CMC_Force_, *t_21_*= -1.23, *p* = 0.23, *d* = 0.01).

### 3.4 Comparison between CMC_EMG_ and CMC_Force_

Figure 4 presents the result of the correlation analysis between global CMC_Force_ and the more classically computed global CMC_EMG_. The magnitude of CMC_EMG_ did not differ significantly from that of CMC_Force_ (*t_35_* = -1.71, *p* = 0.096, *d* = 0.34) and a correlation analysis between the two revealed a strong positive correlation (*ρ* = 0.74, *p* < 0.0001). Of note, removal of the two participants who appear as outliers on Figure 4A did not change the interpretation of the correlation (*ρ* = 0.69, *p* < 0.0001). In terms of CMC significance, although a significantly higher number of participants had a significant CMC_Force_ compared to CMC_EMG_ (*χ*^2^*_1_ =* 9.03*, p* = 0.003), the intercategory association test revealed that participants who had a significant CMC_EMG_ were more likely to also have a significant CMC_Force_ (*κ* = 0.52, *z* = 3.35, *p* = 0.0008; Figure 4B).

**Figure 4.**
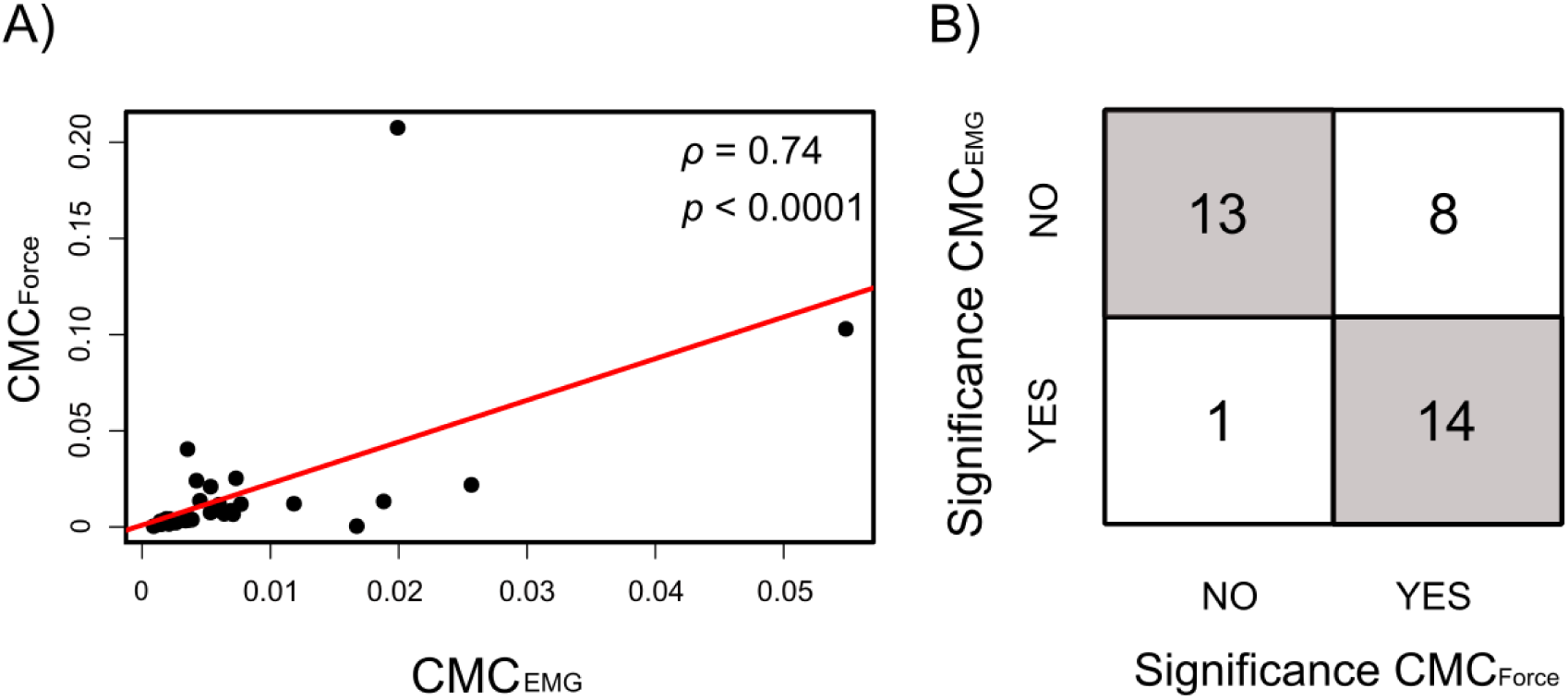
Comparison between CMC_EMG_ and CMC_Force_. A — Correlation between CMC_EMG_ and CMC_Force_. B — Significance confusion matrix for CMC_EMG_ and CMC_Force_. Grey color denotes cells of Significance CMC_EMG_ – Significance CMC_Force_ co-occurrence.

### 3.5 Stability of FDI EMG and force signals during the brachium movement

To verify that the suppressed CMC resulted from the non-focal SM1 beta ERD, and not from brachium movement-induced changes in the FDI EMG and force signals, we investigated the stability of EMG amplitude and force amplitude across T_before_, T_during_, and T_after_. These analyses revealed that, in spite of the presence of movement-induced fluctuation in EMG amplitude in one participant (Figure 5), across all retained participants including the one with movement-induced EMG amplitude fluctuations, there was no significant difference in the average SD of the normalised FDI EMG amplitude between the 3 time periods (*F*_2,42_ = 0.07, *p* = 0.93, *η*^2^ = 0.004). However, there was a significant difference in the SD of the normalised force amplitude between the 3 time periods (*F*_2,63_ = 8.04, *p* < 0.001, *η*^2^ = 0.23), with a greater SD at T_during_ compared to T_before_ (*t*_21_ = 4.73, *p* < 0.001, *d* = 1.12) and T_after_ (*t*_21_ = 3.51, *p* = 0.002, *d* = 0.99). This latter finding was driven by a subset of participants (N = 4; 18 %) in whom brachium movement execution resulted in transient fluctuations in FDI force amplitude (Figure 5). However, across all retained participants including those with movement-induced force amplitude fluctuations, we found no significant correlation between this force signal amplitude variability at T_during_ and the index of movement-related CMC_Force_ modulation across the time periods of interest (*ρ* = -0.11, *p* = 0.64). These results suggest that the observed movement-induced differences in CMC_Force_ (Figure 3B) were not due to this transient increase in force signal amplitude.

**Figure 5.**
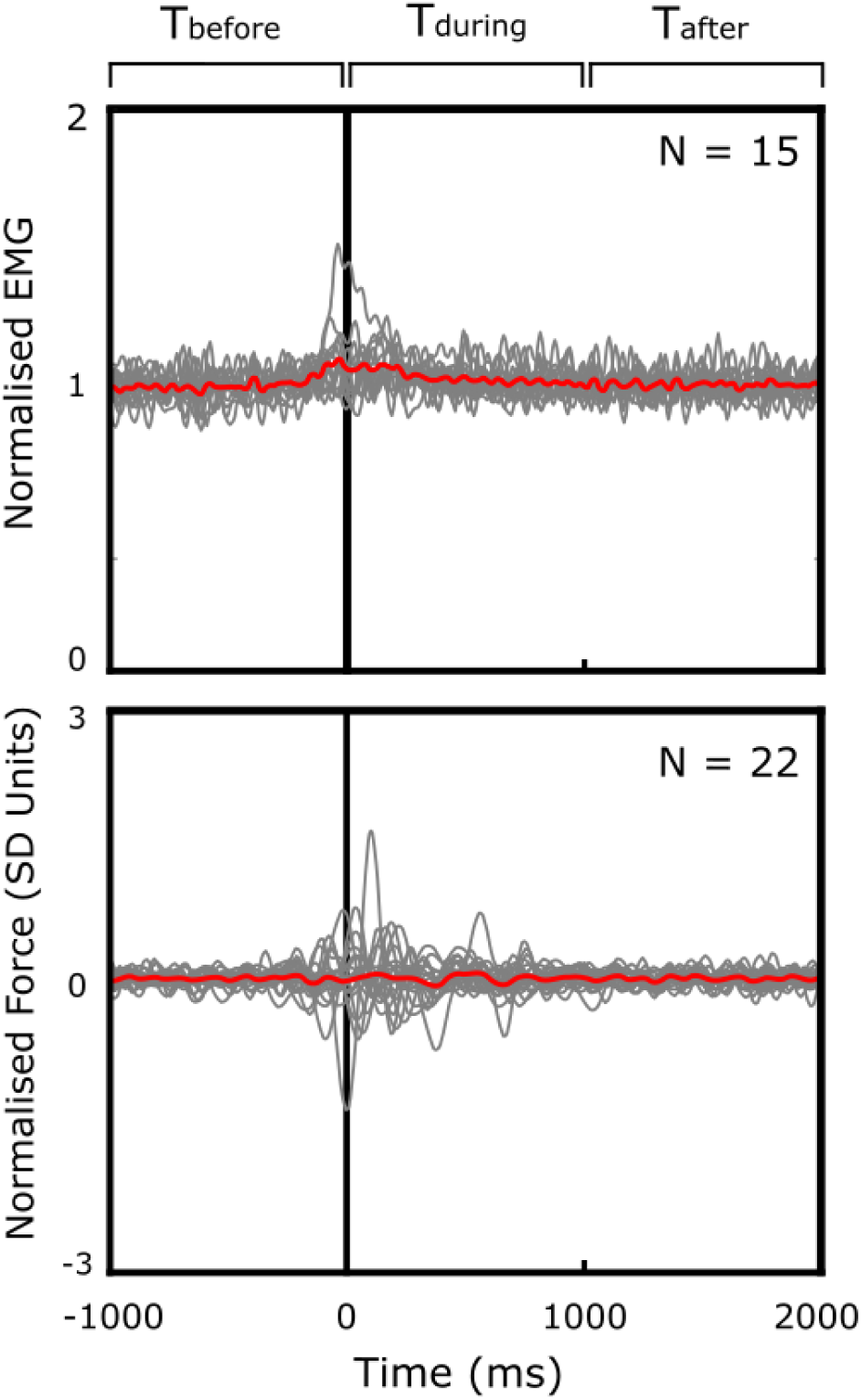
Subject-averaged rectified and normalised FDI EMG (top) and normalised force (bottom) traces for participants with significant CMC. EMG traces were normalised by their mean and the force traces were normalised by their SD. The red lines indicate the grand averaged trace across participants and the black vertical lines indicate the brachium movement onset as identified in accelerometer recordings.

### 3.6 Effect of brachium movement execution on CMC for participants with stable EMG and force signals

To further verify that the observed effect of brachium movement execution on CMC was not due to amplitude instability in the EMG and force signals, we repeated the analyses reported in section 3.2 after excluding the participants with brachium movement–induced transient EMG (N = 1) and force (N = 4) amplitude fluctuations (Figure 5). For CMC_EMG_, the effect of time was still significant (*F*_2,26_ = 4.05, *p* = 0.029, *η*^2^ = 0.237) with CMC_EMG_ magnitude being significantly lower at T_during_ compared to T_after_ (*t_13_*= -2.22, *p* = 0.044, *d* = 0.56). For CMC_Force_ the effect of time was also still significant (*F*_2,42_ = 6.90, *p* = 0.003, *η*^2^ = 0.289) with CMC_Force_ magnitude being significantly lower at T_before_ and T_during_ compared to T_after_ (*t_17_* = -2.52, *p* = 0.022, *d* = 0.40 and *t_17_* = -3.59, *p* = 0.002, *d* = 0.36, respectively). Hence, these results further support the argument that observed movement-induced differences in CMC were not due to transient increase in EMG or force signal amplitude.

## 4. Discussion

In the present work, we used a dual movement execution task involving transient movements of the right brachium and a stationary right FDI muscle contraction to investigate the focality of movement-induced beta amplitude changes along the SM1 homunculus. As classically reported, we observed a contralateral SM1 beta ERD during movement preparation and movement execution. The ERD was followed by a beta ERS after movement termination. These left SM1 beta modulations were accompanied by a suppression and then enhancement, respectively, of CMC magnitude between the left SM1 and the right FDI muscle. Furthermore, similar results were obtained when using CMC derived from EMG and force recordings which further establishes the equivalence of these two measures.

Consistent with a rich literature (Bonaiuto et al., 2021; Georgiev et al., 2025; Nakayashiki et al., 2014; Pfurtscheller, 1981; Salenius & Hari, 2003; Szul et al., 2023; Zaepffel et al., 2013), brachium movement execution resulted in a significant beta ERD in the contralateral SM1 shortly before and during movement execution, which was followed by a significant beta ERS after movement termination. The rather large temporal span of the observed beta ERD likely reflects a combination of top-down anticipatory attention during the expectation of the auditory tone, preparatory changes in SM1 excitability at tone perception, and the actual activation of the SM1 resulting in motor efference after the tone (Khanna & Carmena, 2017). Importantly, the CMC with the FDI muscle followed a very similar pattern of modulations with an attenuated CMC magnitude observed shortly before and during movement execution and an enhanced CMC magnitude observed after movement termination. These CMC magnitude changes were not accompanied by changes in CMC peak frequency, which, according to some views, indicates steady corticomuscular communication in the absence of error-correcting motor drive (Mehrkanoon et al., 2014).

Following the rationale outlined in the introduction, our finding that CMC with the FDI underwent similar modulations as the SM1 beta amplitude favours the hypothesis that beta amplitude modulations are non-focal. Nevertheless, the amplitude of brachium movement-induced ERD did not show a significant impact on CMC magnitude, as indicated by the absence of significant differences between the CMC magnitude during movement execution trials with low compared to high beta ERD. It could be expected that CMC magnitude would show its greatest decrease when beta ERD is high (and hence beta amplitude is the most suppressed), however, our results were not able to capture this effect likely due to insufficient number of trials per participant. Given that a well-known determinant of CMC magnitude is the presence of beta oscillations in a given SM1 area and its target muscle (Bourguignon et al., 2017; Bräcklein et al., 2022; Echeverria-Altuna et al., 2022; Mongold et al., 2022), it is reasonable to assume that, in the present study, the suppressed CMC during movement execution resulted from a spread of beta ERD from the SM1 brachium area to the SM1 FDI area. Given that CMC is mediated by beta corticomuscular transmission through only the fastest corticospinal fibers (Ibáñez et al., 2021), influences on CMC other than spreading beta ERD, such as for instance phase non-stationarity of beta oscillations, are unlikely to affect CMC magnitude. Any such non-stationarity at the cortical level would be propagated to the peripheral signals in a time that is far too short (∼20 ms; Ibáñez et al., 2021) to affect the CMC estimation.

This argument is in line with previous findings of invasive electrophysiology in animals that has established functionally relevant propagatory effects of beta oscillations along the SM1 (Balasubramanian et al., 2020; Best et al., 2017). It is also in line with electrocorticography recordings in humans that have highlighted wide patterns of movement-related beta suppression (Miller et al., 2010) as well as beta propagatory effects in the SM1 (Stolk et al., 2019). Moreover, our finding further supports the results of Zich et al. (2023) who identified similar movement-induced beta propagatory dynamics with MEG. Considering the anatomical organization of the SM1, our results likely capture mostly a medial-lateral (i.e., parallel to the pre- and postcentral gyri) beta propagation reported by Zich et al. (2023).

Our finding that beta ERD induced by the brachium movement affects the CMC between the SM1 FDI area and the FDI muscle, even though the FDI muscle was stationary and not involved in the movement execution, raises important questions for the role of beta oscillations in motor control. Part of the explanation may lie in the tendency of body effectors to be coupled at the biomechanical and functional level. At the biomechanical level, due to reaction forces, executing a movement of one body segment entails anticipatory and compensatory adjustments of muscles acting across other kinematically-linked segments (Buhrmann & Di Paolo, 2014; Maeda et al., 2017). In this context, propagated beta ERD/ERS could reflect the necessary engagement of the SM1 areas in control of these later muscles. At the functional level, bodily segments are seldom moved in isolation. Instead, purposive actions entail coordinated activation of multiple muscles, which are believed to be encoded in an action repertoire in the primary motor cortex (Graziano, 2006). Accordingly, moving a body segment is likely to activate several of the encoded patterns, and brain structures like basal ganglia and the cerebellum help ensure that the relevant motor command is being sent down the corticospinal tract. Collectively, these considerations suggest that non-focal SM1 cortical beta ERD and ERS reflect regulatory mechanisms that coordinate the coactivation of multiple muscles and assure the precision of arm movement.

The finding that CMC computed with the force signal of a contracted muscle (CMC_Force_) is strongly related to the CMC computed with the EMG of that muscle (CMC_EMG_) and that the two show very similar patterns of movement-induced modulation further establishes force recordings as a valid surrogate for EMG for CMC estimation (Airaksinen et al. 2015; Bourguignon et al. 2017). Here, we build upon previous MEG findings and show that force-based CMC estimation can also be reliably performed with EEG. However, it is worth keeping in mind that EEG has a lower signal-to-noise ratio than MEG (Destoky et al. 2019), which accounts for the lower proportion of participants with significant CMC as well as the overall lower coherence values in our study compared to the previous MEG investigations (Airaksinen et al. 2015; Bourguignon et al. 2017). Moreover, we observed a significantly higher percentage of participants having a significant force-based CMC compared to EMG-based CMC. This suggests that peripherally-propagated beta bursts are more clearly visible in the force signal compared with the EMG signal where the beta bursts would be embedded within the ongoing muscle activity. Collectively, these considerations suggest that EEG-force CMC can provide an accessible, yet equally if not more resolute, way to assess SM1 beta oscillations’ propagation towards the periphery. This can be particularly useful in investigations with clinical populations such as stroke or Parkinson’s disease where abnormalities in CMC have been associated with impaired motor behavior (von Carlowitz-Ghori et al., 2014; Zokaei et al., 2021). Furthermore, leveraging CMC as a proxy for characterizing the focality of beta oscillatory activity along SM1 homunculus can allow for non-invasive assessments of the topographical spread of those oscillations, which is known to be affected in, for instance, Parkinson’s disease (Moisello et al., 2015; Nelson et al., 2017).

Although we did not find a significant difference in the FDI EMG signal stability before, during, and after the brachium movement, we did observe some participants for whom the brachium movement execution resulted in transient fluctuations in the force. Similarly, force fluctuations were also identified in a study assessing the impact of auditory and visual distracting stimuli on CMC (Piitulainen et al., 2015). In this study, the force fluctuations were hypothesized to result from reflexogenic mechanisms activated by a startle-like response to the external stimulations (Piitulainen et al., 2015). It is therefore possible that the transient increase of the force we observed in some participants in our study is a representation of this startle-like effect caused by the auditory tone that instructed our participants to initiate the movement execution. Alternatively, it could also result from changes in FDI SM1 efference in anticipation and compensation of the brachium movement execution itself. In any case, these force instabilities did not affect the presented results and did not explain the observed CMC modulations.

Noteworthily, a limitation of our approach is that, while CMC hinges on the presence of the beta oscillations within a given SM1 area (Bourguignon et al., 2017), it assesses phase and amplitude dependent corticomuscular coupling, and does not directly assess local beta amplitude in that given area. Therefore, future work with invasive electrophysiology is needed to fully validate the approach we propose whereby CMC is used to assess the focality of beta ERD.

A further limitation is that the two SM1 regions investigated in our study, the SM1 brachium area and the SM1 FDI area, are located in close anatomical proximity to each other. Therefore, based on our results, we cannot establish the influence of the spreading beta ERD along the whole SM1 homunculus. Zich et al. (2023) identified beta propagation within a distance of approximately 6 cm^2^ and our results are consistent with that finding as the brachium and FDI areas are within that distance. However, future studies including dual tasks involving pairs of motor effectors at varying anatomical distance are warranted to further characterize the degree of non-focality of the beta ERD.

An additional limitation of our work is the reduction in effective sample size resulting from the selection only of participants with significant CMC. Although this is a common practice in CMC studies (e.g., Murnaghan et al., 2014; Perez et al., 2012; Piitulainen et al., 2015; Rossiter et al., 2012; Sharifi et al., 2021; Suzuki & Ushiyama, 2020), it implies that the effects we observed may be present only in the participants with the greatest CMC, or in other words, with the most efficient transfer of cortical beta oscillations towards the muscles (Bourguignon et al., 2017). Future replications of our results with MEG are likely to improve upon this limitation as the superior spatial resolution and signal-to-noise ratio of MEG (Destoky et al. 2019) are expected to enhance the detection of both cortical beta oscillations and their coupling with muscle activity.

In conclusion, our results show that movement-induced beta ERD in one SM1 area alters the CMC between the SM1 and a muscle not involved in the movement execution. This suggests that the beta ERD could spread along the SM1 homunculus and induce a suppression of beta oscillations in neighbouring areas as well. Importantly, the combination of a dual motor task and CMC evaluation to assess this spreading non-invasively is novel, and could be of interest for the appraisal of clinical populations where beta oscillations are altered such as in stroke or Parkinson’s disease. We further demonstrate that CMC suppression can be captured not only via EMG-based but also through force-based recordings. This establishes force-based CMC as a viable technique that can make investigations of SM1 beta oscillations more broadly accessible for healthy and clinical populations.

## Author Contribution

**CG**: investigation, data curation, formal analysis, visualization, writing – original draft; **SJM**: investigation, data curation, writing – review and editing; **GN**: methodology, supervision, writing – review and editing; **MB**: conceptualization, data curation, formal analysis, methodology, supervision, funding acquisition, writing – review and editing

## Data availability statement

All anonymized data is available on the OSF (https://osf.io/hcb24/overview).

## Acknowledgements

CG and SJM were supported by the FNRS (Brussels, Belgium). GN is a postdoctoral specialist at the FNRS. MB was supported by the FNRS, the Brussels-Wallonia Federation (Collective Research Initiatives grant), and the Walloon Region strategic axis (FRFS-WELBIO).

## List of Abbreviations

ANOVA: analysis of variance
ACC: accelerometry
CMC: corticomuscular coherence
CMC_EMG_: corticomuscular coherence computed with EMG signals
CMC_Force_: corticomuscular coherence computed with force signals
EEG: electroencephalography
EMG: electromyography
ERD: Event-Related Desynchronization
ERS: Event-Related Synchronization
FDI: first dorsal interosseous
MEG: magnetoencephalography
MVC: maximum voluntary contraction
SM1: primary sensorimotor cortex
SD: standard deviation
T_before_: 1000 millisecond time period before movement
T_during_: 1000 millisecond time period during movement
T_after_: 1000 millisecond time period after movement
TFR: time-frequency representation

